# TomoScore: A Neural Network Approach for Quality Assessment of Cellular cryoET

**DOI:** 10.1101/2024.11.06.622356

**Authors:** Xuqian Tan, Xueting Zhou, Ethan Boniuk, Anisha Abraham, Zhili Yu, Valerie Dalton, Steven J. Ludtke, Zhao Wang

## Abstract

Electron cryo-tomography (cryoET) is a powerful imaging tool that allows three-dimensional visualization of subcellular and molecular architecture without chemical fixation. Tomogram quality varies widely, particularly during large high-throughput data collections, and the most common strategy for initial quality assessment is empirical judgment by an expert. Tomograms are collected for two distinct purposes: 1) Annotation of subcellular features and cellular morphology, typically performed at lower magnifications and higher defocus. 2) Subtomogram averaging, generally at higher magnifications and closer to focus. For the first purpose, contrast and the ability to distinguish cellular features of interest are key, whereas for subtomogram averaging, recoverable signal at high resolution is the key factor. We have developed “TomoScore” a deep-learning based tomogram quality metric to assess the suitability of individual tomograms for cellular annotation based on the scale of features which can be distinguished.

We further explore the relationship between accumulated electron dose and resulting quality, suggesting an optimum dose range for cryoET data collection. Overall, our study streamlines data processing and reduces the need for human involvement during pre-selection for tomogram segmentation.

## Background

Electron cryo-tomography (cryoET) is currently the only technique capable of providing 3D structure of unlabeled/unfixed cells at nanometer resolutions^1,2^. This method is also emerging as a technique for subtomogram averaging, providing high-resolution macromolecular structures *in-situ*^3^. While these two goals seem related, they generally require significantly different data collection parameters, and thus there is relatively little overlap in data sets for the two purposes. Since subtomogram averaging requires fine pixel sampling to achieve high resolution averages, the field of view is dramatically restricted making them much less suitable for studies of subcellular organization. For cellular tomogram annotation, lower magnifications provide larger fields of view, and higher defocus provides significantly improved low-resolution contrast to increase annotation accuracy. While subtomogram averaging tools, such as CTFFIND^4^, provide good estimates of whether tilt series contain enough high-resolution information to be suitable for that task, CTF analysis fails to capture useful information about suitability for cellular morphological annotation. Fourier contrast is often present even when tomograms lack sufficient features for annotation.

Accurate interpretation of cellular tomograms requires annotation of features at different levels of detail, ranging from individual macromolecules to organelles depending on the purpose of the study^5–7^. However, tomogram quality can vary widely due to imaging conditions and how the cells were prepared for imaging. For example, a thicker lamella provides a larger cellular volume but decreases the contrast of individual features within the cell. When a project involves hundreds to thousands of cellular tomograms, prescreening tomograms to assess suitability for annotation is critical. This process is often performed manually by a human visually screening and, in some cases, trying to annotate each tomogram in the set^8,9^. To reduce the need for this time-consuming process and to reduce potential human bias, we have developed a deep-learning based quantitative quality metric that can help rank tomograms. The “quality” we assess in this context is a measure of the smallest reliably resolvable feature present in individual 2D slices of the tomogram. The quality metric is determined for each tomogram slice, providing both an overall quality metric and a thickness estimate(see more details in methods). This metric is completely independent of typical resolution measures, such as the FSC between tomograms reconstructed from even movie frames and odd movie frames, which can produce both over- and underestimates of the tomogram quality for annotation purposes^10–13^. Therefore, our metric only accesses the quality of the final cellular reconstructed tomograms in their “as-is” state.

Since the problem of scoring tomograms is mathematically ill-posed, we made use of deep learning techniques, building on ResNet^14^, with human-derived training data to produce our trained SliceQuality Model. The network was trained across multiple cellular species, microscopes, and instrument settings. The strategy we used for manually quantifying quality is straightforward and has minimal annotator-dependence.

## Results

### Human-assessed quality categories for network training

Our goal is to produce a quantitative measure of tomogram quality mimicking human judgement. Since human judgement on such a task lacks precision and consistence, we opted to limit human judgement to 6 discrete categories for network training. With significant amounts of training data, the final trained network can produce a continuous quality value spanning the original categorical space. The categories identify the size in pixels of the smallest reliably human-discernible biological feature present in each slice (Fig. 1, Fig. S1). In addition to the 5 discrete feature-size categories presented in Fig. 1, a sixth category is reserved for tomogram slices with no discernible biological features at all. This simple per-slice categorization can be rapidly assessed, allowing us to generate a sufficiently large training set without expending a prohibitive amount of human effort.

**Figure 1.**
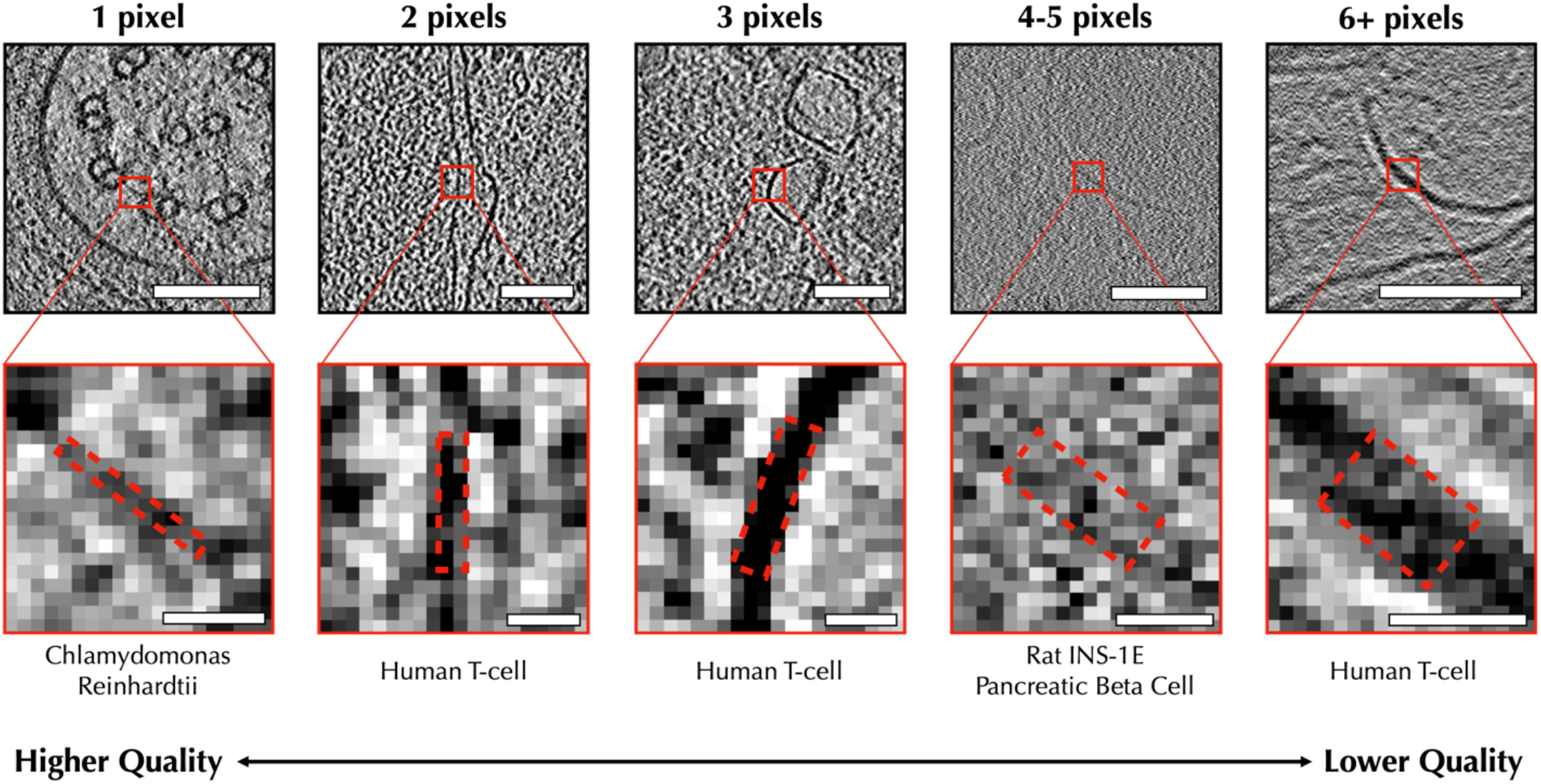
Demonstration of different quality tomogram slices under human-defined scoring criteria. This figure demonstrates how membrane features are scored based on their size in pixels. Example features are included from C. reinhardtii (left), Human T-cells (mid-left, middle, right), and Rat INS-1E pancreatic beta cells(mid-right). The dashed red rectangles indicate the width measured for each feature. Images on the top row have been cropped to 200×200 pixels with a scale bar of 100 nm. Images on the bottom row have been cropped to 20×20 pixels with a scale bar of 10 nm. From left to right, the angstrom per pixel values of each image are 13.68, 18.92, 18.92, 14.08, and 9.867 Å/pix respectively. Additional examples taken from other species, as well as measurements for particle features, can be found in Supplementary Fig. 1.

Due to the resolution anisotropy of tomograms, annotation is typically performed on 2-D slices in the x-y plane where resolution is fairly isotropic, so our quality metric is similarly implemented slice-wise. As this is a quality metric and isn’t performing actual annotation, incorporating Z information would add little to the analysis, and slice-wise analysis provides the side-effect of producing a thickness estimate. To build a training set spanning commonly observed cellular features, we selected various prokaryotic and eukaryotic cells from the Cryo-ET Data Portal (Chan Zuckerberg Imaging Institute, Chan Zuckerberg Initiative)^15^. We also included platelet tomograms generated by our lab as a specialized type of eukaryotic cell (anucleate) due to their unique cellular structure (Supplementary Table 1). Overall, 58,527 slices from 114 tomograms across 19 species were gathered and manually assessed.

During the human assessment of tomogram quality, we observed a logical trend: within one tomogram, central slices tend to have the highest quality, while the top and bottom slices have much lower quality. To encourage training the network to produce a smooth gradation between discrete scores, we took the per-slice discrete quality values along Z and smoothed them with a Gaussian filter with z=3 pixels.

We employed the following two strategies during data partitioning to establish rigorous training. First, we performed a tomogram-based partition for unbiased training. Specifically, 98 of 114 tomograms were randomly selected for the training set pool, while the remaining tomograms were left for validation (1/114) and testing (15/114). Next, we performed a sample size partition for balanced training. Among the selected 98 tomograms, 2500 slices were randomly selected from each human labeled category to be added to the training, ensuring the prediction is not affected by sample size. Our final training set contained a total of 15,000 slices evenly distributed across six categories with corresponding continuous quality estimations.

### SliceQuality Model, a modified ResNet101 network, provides per-slice quality estimation for a wide range of cyroET samples

We adopted the structure of ResNet101^14^ and modified it to predict the quality score of a single tomogram slice, as shown in Fig. 2a. Specifically, to obtain an output quality score ranging from 0 to 1, we added a sigmoid function at the end of ResNet101. Our continuous quality estimation in the training set uniformly map to a 0-1 prediction range (0: no biological feature; 0.2: 6+ pixels; 0.4: 4-5 pixels; 0.6: 3 pixels; 0.8: 2 pixels; 1.0: 1 pixel). Our model takes less than 30 seconds to produce predictions for a standard 1k*1k tomogram containing ∼200 slices on a machine with an NVIDIA GeForce RTX4090 GPU. A speed that substantially exceeds manual labeling.

**Figure 2.**
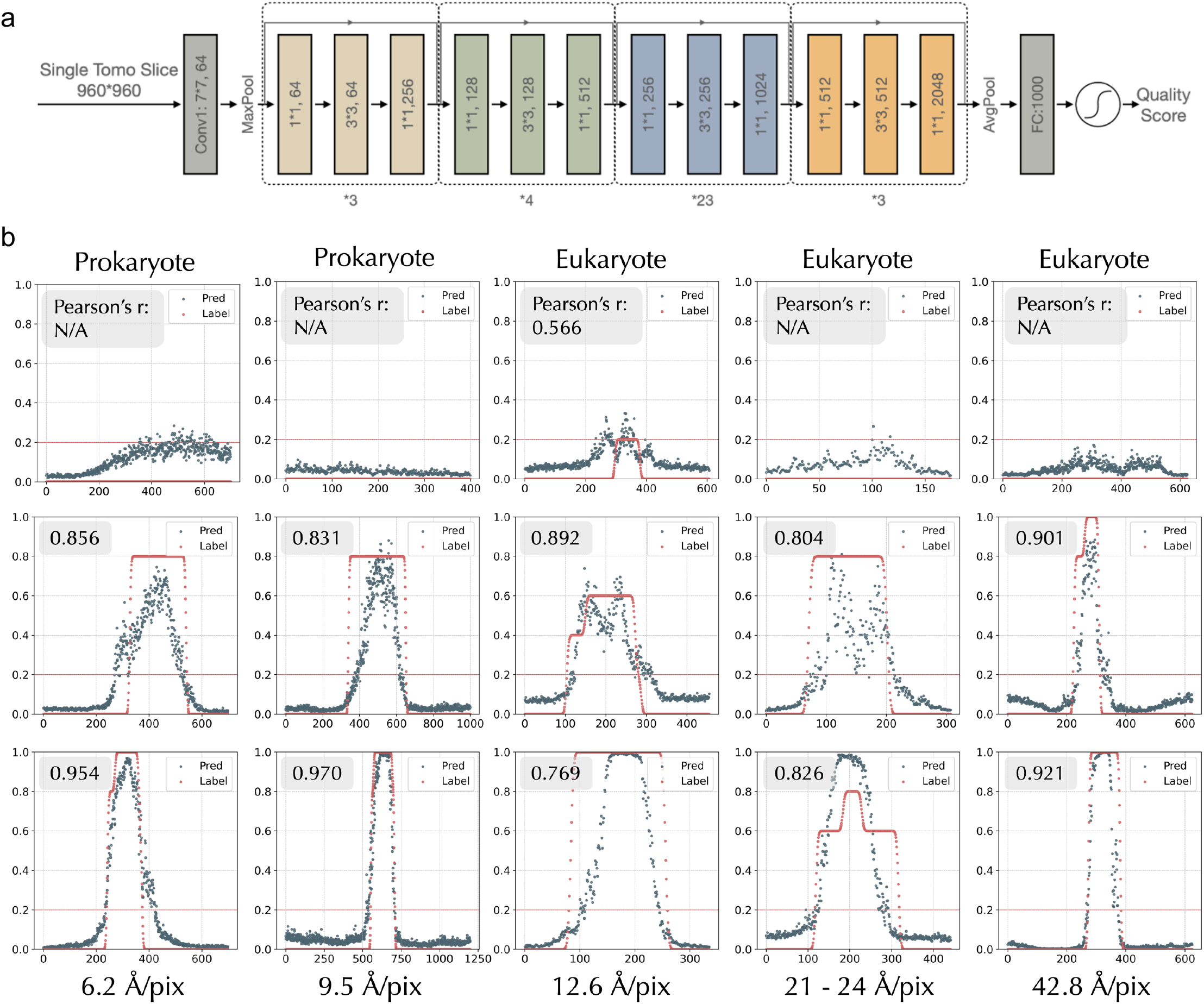
SliceQuality Network structure and training result. a) SliceQuality Network is modified from ResNet101 structure. A sigmoid function is appended before the output layer to give a predicted quality score within the range 0-1. b). SliceQuality Network’s prediction score vs. label based on slice position curve for 15 test tomograms excluded from the training process. Pearson correlation coefficient is calculated (labeled on the top left) for each tomogram’s prediction-label pair, and all of them are statistically significant (p<0.001). Four of the lower-quality tomograms did not receive the Pearson correlation coefficient due to their constant labeling of 0. Each image has been cropped to 960×960 pixels for comparison. The first row captures the following, from left to right: *Sulfolobus solfataricus, Treponema primit*ia, human umbilical vein endothelial cell (HUVEC), human blood platelet, and human blood platelet. The middle row captures the following, from left to right: *S. solfataricus, Vibrio cholerae*, HUVEC, HUVEC, human blood platelet. The last row captures the following, from left to right: *Magnetospirillum magneticum, V. cholerae*, HUVEC, HUVEC, and human blood platelet.

To test model accuracy and generalizability, we selected 15 tomograms as the test set across different cellular species, microscopes, and magnifications. To quantitatively measure model accuracy, we overlaid the label scores and predictions and then plotted them by each slice’s position within one tomogram. An average Pearson correlation coefficient of 0.845 (p < 0.001) is reached between the prediction and label (Fig. 2b). Notably, our model also performs well on the three species (*Sulfolobus solfataricus, Treponema primitia, and Magnetospirillum magneticum*) that were not included in the training set, demonstrating its generalizability. We also noticed that two of the tomograms (row2, col2, and row3, col3) have a wider range of higher-quality slices for human labeling. After manual inspection, we found that both tomograms had tilted samples, resulting in the clear-cellular feature proportion of top and bottom slices being smaller than usual, thus receiving a lower predicted quality score. Further, our model also demonstrated its ability to differentiate between biological features and non-biological features, such as empty holes (row1, col3) and ice layers (row1, col4).

### SliceQuality Model accuracy and consistency were further validated by tomogram accumulated dose *in silicon* variations

The question of what dose to use when collecting tilt series has two independent components. First, the dose required to produce sufficient contrast in the tomogram to test the experimental hypothesis and second, at what dose level radiation damage begins to distort results. We were able to separate these variables via two experiments. First, we collected a tilt series with high cumulative dose (400 e^-^/Å^2^), collecting each tilt as a movie to provide dose fractionation. We then produced tilt series using only a fraction of the frames at each tilt to generate tilt series with lower dose (from a noise perspective) but near-identical radiation damage. These tilt series allow us to investigate how tomogram quality varies with dose in a damage-independent manner. Second, in a new region we collected a sequence of tilt series, each with low cumulative dose, on the same region. As demonstrated in later results, each of these tilt series has the same dose from a noise persepective, but each begins with the cumulative radiation damage of the previous series. This allows us to monitor the impact of radiation damage on the measured tomogram quality

Assuming that our model is consistent and accurate, two identical tomograms should produce the same quality score using our model. However, practically, identical tomograms do not exist. Therefore, we developed an *in silico* method to create “identical” tomograms to test model consistency and accuracy (Fig. 3a). First, we collected a separate dataset containing 10 platelet tomograms with a high cumulative dose. Specifically, we recorded 152 raw frames per tilt image, with the total dose per tilt series being 400 e^-^/Å^2^. The movie frames from each angle were separated into even/odd stacks. The only variation between these stacks should be noise since the aggregate dose at each even/odd movie frame differs only by a tiny amount. The even/odd tilt series were reconstructed using the same tilt geometry and other parameters (Fig. 3b). When the FRC is computed between even/odd slices of these reconstructions there is minimal variation throughout the reconstructed thickness. That is, per-slice FRC is unable to detect whether features suitable for annotation are present in any given Z-slice. Our model, on the other hand, can clearly distinguish the quality differences along Z.

**Figure 3.**
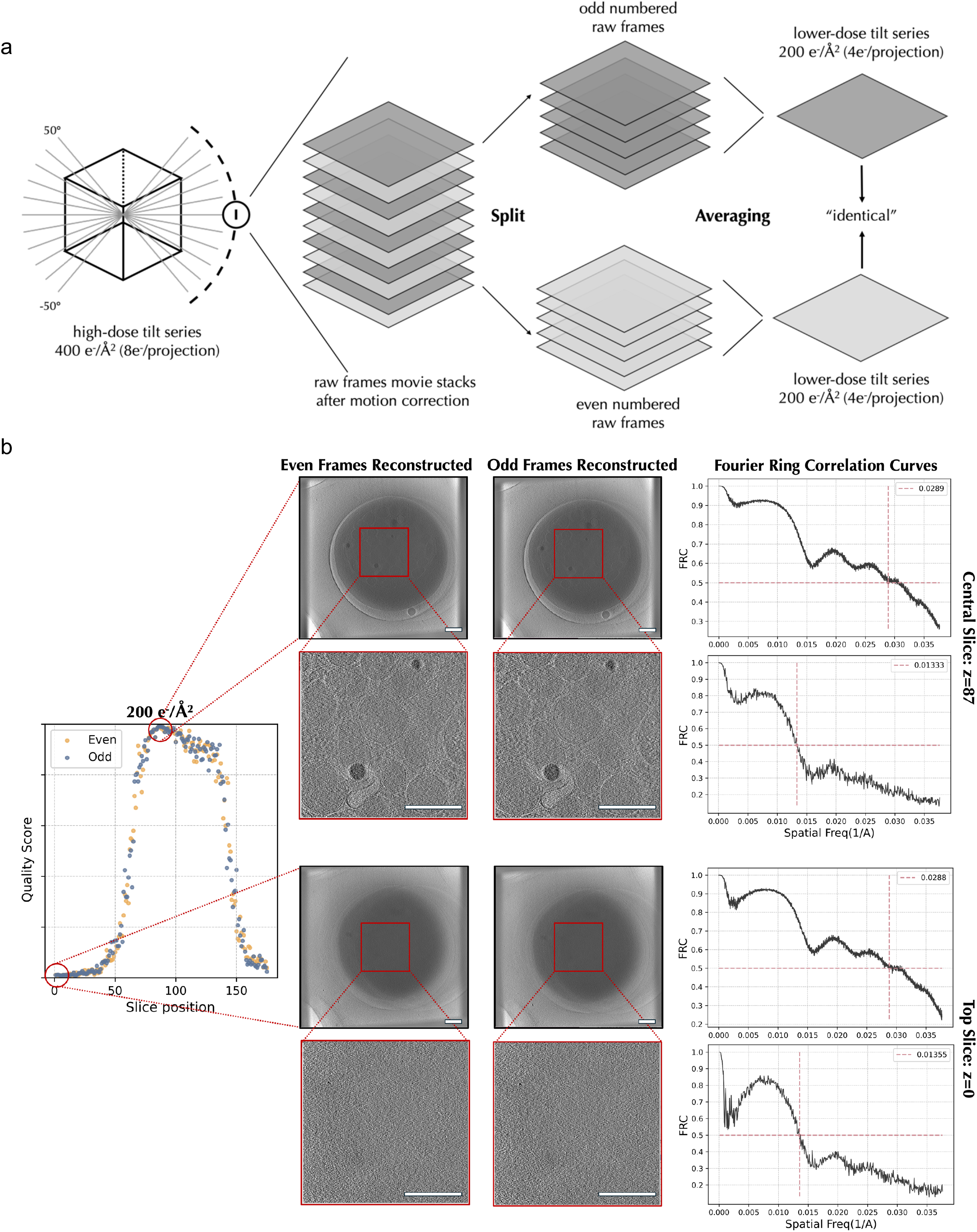
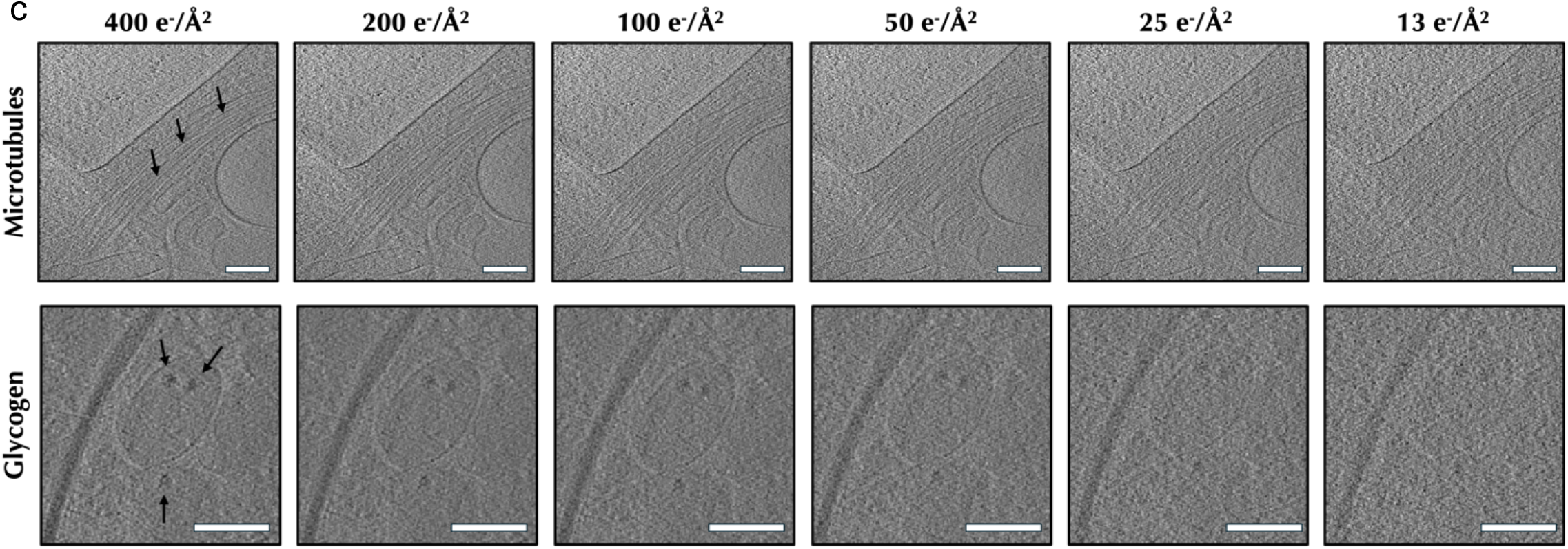
Model consistency tests on generated “identical” tomograms. a) A workflow of separating even/odd raw frames from a single 2D tilt projection. Separated raw frames were averaged and merged into two parallel 2D tilt projections. Repeat this process for all tilt projections in a series to generate two identical sets of tilt series such that each set is exactly half dose of the original tilt series. Each set of half-dose tilts was then reconstructed into a 3D tomogram using the same alignment parameters. b) The even/odd frames reconstructed tomogram slices are similar by visual comparison. The Fourier ring correlation (FRC) curve also shows that both central and top slices from the two tomograms share high similarities at lower resolution frequencies. On the right, quality scores predicted by our trained model indicate that central slices have better resolvability (i.e higher quality score), where the FRC curves fail to show. c) Level of radiation exposure differentially affects tomogram quality. At higher radiation doses, tomograms of the same platelet exhibit higher-quality structures of microtubules and glycogen. At lower doses, tomograms exhibit more noise and structures are less defined. Scale bars are 200 mm

We then generated a series of tomograms with different cumulative doses (13 e^-^/Å^2^ to 200 e^-^/Å^2^) from the same tilt series by averaging different proportions of the movie frames from micrographs (see method section for details). Using our SliceQuality Model, we obtained the per-slice quality scores for the tomogram series with different accumulated doses, shown in (Supplementary Fig. 2a). We further confirmed consistency and robustness between tomogram score predictions generated from even/odd frames at varied accumulated dose levels. The consistency between even/odd reconstruction predicted qualities decreases in data with a lower cumulated dose. Notably, all even/odd quality score differences were significantly lower than 16.7% (⅙), which is the discrete step size of the human categorization. This result indicates the prediction is on par with human judgments (Supplementary Fig. 2b). Thus, we confirmed that our SliceQuality Model generates robust quality predictions at different doses.

Furthermore, we observed that the predicted quality score peaks decrease as the accumulated dose decreases, indicating that there’s a decrease in overall tomogram quality along with a decrease in the simulated cumulative dose used in the reconstruction^16^. We performed a visual comparison at the same position in the tomogram to validate the changes in structural feature resolvability according to the dosage. We chose two specific positions within the tomograms that effectively encapsulate the key features considered in the initial ranking criteria. As shown in Fig. 3d, 400 e^-^/Å^2^, 200 e^-^ /Å^2^, and 100 e^-^/Å^2^ tomograms clearly show microtubules with linear morphology and separation between neighboring objects allowing for clear segmentation. Glycogens appear as high-contrast circular structures with precise boundaries to the cell environment in 400 e^-^/Å^2^ and 200 e^-^/Å^2^ tomograms. At lower dosages, organelles such as mitochondria lose contrast, which compromises the ability to visually identify the organelle by distinct features such as the double membrane and cristae. These results and observations inspired us to come up with the following metrics.

### A novel metric, TomoScore, was created for cryoET quality assessment

As we previously defined, a predicted quality score of 0.2 maps to the threshold for the presence of biological features (6+ pixel size). We consider any slices scored above 0.2 to contain biological features that could be annotated. Therefore, we estimate the sample thickness by finding the range of slices scoring above 0.2. We validated the estimated thickness with manual thickness measurements from YZ projections (Supplementary Fig. 3a,b,c) on 137 tomograms (excluded from the training) through the RANSAC algorithm^17^ (Supplementary Fig. 3d) and obtained a regression slope of 0.988 (p<0.001). We manually examined each outlier and concluded three causes led to inaccurate thickness measurements: 1) The missing regions between angular steps in the tilt series generally lead to streaks from high contrast features in the tomograms, which can propagate beyond the thickness of the actual specimen. 2) The reconstruction may be modestly tilted with respect to the X-Y plane, leading to “good” slices extending over a larger range of thicknesses. 3) Low tomogram resolvability.

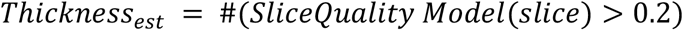

To simplify the inter-tomogram quality comparison, we created a metric called TomoScore. TomoScore is generated from the predicted quality scores of a single tomogram and can be interpreted as the combination of all slice quality scores normalized by the estimated specimen thickness. As a result, TomoScore calculates the average quality score of slices within a single cell to represent the overall quality of a tomogram. Consequently, TomoScore has an expected range between 0 and 1, where 0 indicates no slice within the tomogram reveals cellular structures and 1 would denote a theoretically perfect tomogram with all slices revealing the optimal quality of organelle structures.

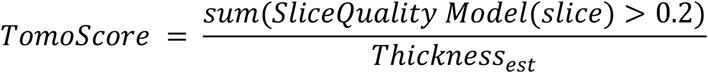

We further validated our model’s consistency on TomoScore using the same even/odd frames split tomograms. For a single set of tomograms split into different total doses, the TomoScore differences between even/odd tomograms are small across the total dose from 13 e^-^/Å^2^ to 200 e^-^/Å^2^ (Fig. 4a). We also have a small range of distribution for TomoScore differences between the 10 even/odd split tomograms across different total doses (Fig. 4b). To verify that TomoScore is sensitive to dose-induced quality degradation, we performed a control experiment in *E. coli* by repeatedly exposing the same area to 50 e^-^/Å^2^, thereby increasing the cumulative dose while monitoring TomoScore changes (Supplementary Fig. 3). The results show that TomoScore declines as bubbles emerge from high accumulated dose applied to the *E. coli*. This verifies that platelets under lower magnification are less susceptible to visible dose damage even under 400 e^-^/Å^2^.

**Figure 4.**
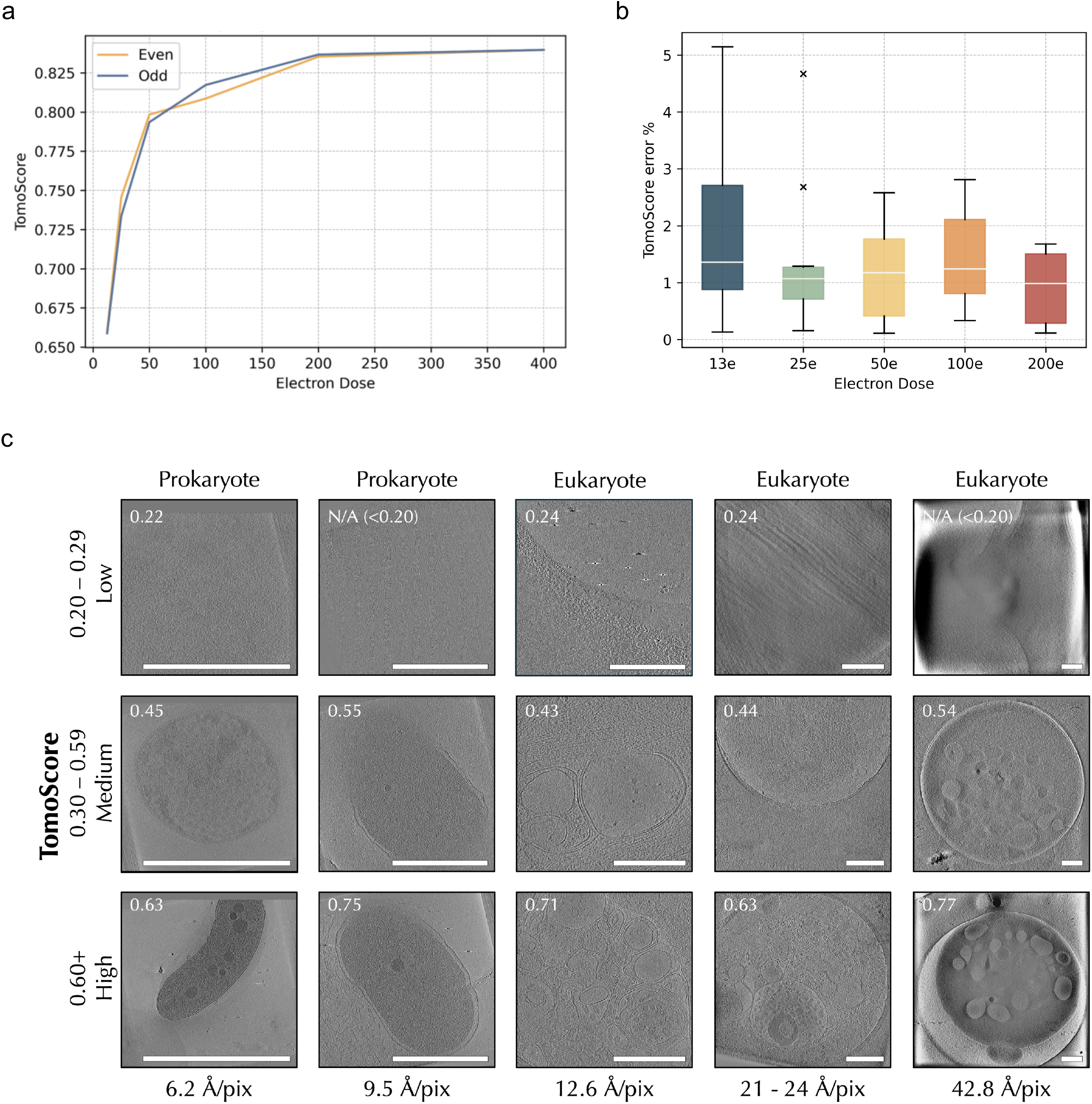
The trained model produces consistent TomoScore. a) Calculated TomoScores for recursive even/odd dose splits of a sample high-dose tomogram. TomoScore decreases as the total doses decrease. b) TomoScore differences distribution for all 10 high-dose tomograms across different split dosages. c) TomoScore accurately predicts tomogram quality across various cell types and magnifications. We compare the most resolvable slice, as determined by a human, of the same 15 tomograms used for Fig. 2b. As predicted, tomograms with a high TomoScore are of better quality and have more resolvable subcellular structures, and all scale bars are 500 nm.

To verify TomoScore aligns with human quality assessment, we visually examined the same 15 tomograms from the test set. Fig. 4c demonstrates how higher TomoScores correspond to higher quality tomogram slices as perceived by the human eye. This trend is preserved between different cell types and magnifications, indicating the broad utility of TomoScore in predicting tomogram quality.

## Discussion

As CryoET has become more automated, large numbers of tilt series can be collected and automatically reconstructed in a single day. Manually assessing the quality of each tomogram in a set of hundreds of tomograms can be a time-consuming process. Having an automated quantitative assessment of tomogram quality both reduces the potential for human bias which might be associated with a given hypothesis and allows subsequent annotation and analysis to focus on the best quality data.

Deep learning techniques have opened new avenues for addressing the challenges of tomogram data processing. Our TomoScore quality evaluation system is especially useful for cryoET, where the datasets are large, complex, and noisy. New data collection strategies, such as Parallel/montage cryoET^18–21^ on Thermo Scientific EPU Software or similar SerialEM methods, can collect multiple locations in one single angular step, increasing the throughput of cryoET data collection compared to traditional collection schemes. Moreover, rapid tilt-series acquisition^22^ or newer continuous tilt methods can acquire a complete tilt series in ∼2 minutes, which further increases cryoET collection speeds. Batch tomogram^23–25^ and most tomogram analysis software packages like IMOD^26,27^, EMAN2^28,29^, Tomoauto^30^, emClarity^31^, Aretomo^32^, and Warp^33^ can run automated or semi-automated reconstruction of tomograms in a pipeline. These improvements have dramatically increased the throughput of cryoET data collection and reconstruction. Therefore, an automated quality assessment tool specific to cellular cryoET is a critical step in high-throughput cryoET analysis pipelines.

In summary, our SliceQuality Model has proven capable of per-slice quality analysis of cellular cryoET across different species and magnifications. The resulting tool demonstrates the ability to accurately, consistently, and, more importantly, quickly assess the resolvability and thickness of reconstructed tomograms. TomoScore thus permits a large set of tomograms from a data collection to be quality-ranked for further analysis. TomoScore is on par with human judgment and is fully automated, saving human effort for downstream processing.

Using TomoScore, we identified the optimal dose range for platelets under a specific set of experimental conditions. Clearly, other variables play a role in this optimal dose, such as properties of the cell type, selected defocus range, lamella thickness, and other issues, but for a given set of conditions, TomoScore does provide a method for optimizing the targeted dose^34–37^. By using an *in silico* approach to control the accumulated dose, the resulting resolvability differences across different doses should not be affected by radiation damage. In the parallel experiment, where we repeat imaging on the same cell, we observed a dose-induced quality degradation, suggest this tool is capable of providing a sensitive measurement of the quality of cellular tomograms sensitive to radiation damage.

Overall, our SliceQuality Model, along with TomoScore can provide an overall estimation of cellular cryoET quality, taking all possible influencers into account. We successfully demonstrated that this method is robust for tomogram quality prediction with high accuracy and consistency. We are confident that our TomoScore can contribute to quality standardization in the cryoET community.

## Methods

### Platelet sample preparation and freezing

Freshly drawn healthy human blood (Gulf Coast Regional Blood Center) was centrifuged at 200G to obtain platelet-rich plasma (PRP) and further centrifuged at 100G to remove the residual red blood cells. We applied 2-4µL PRP to glow-discharged Quantifoil grids and used a Vitrobot (Mark IV, FEI Corp) to quickly freeze samples to form vitreous ice. Mouse platelet samples were taken from previous research^7^.

### Platelet data collection, processing, and reconstruction

Platelet data were collected using a 300 kV FEI Titan Krios microscope with a Gatan K2 Summit direct electron detector camera, through the SerialEM software. The magnification was 11,500x, with a pixel size of 13.26 Å. The tilt series were collected using a 2° angular step, unidirectionally from −50° to +50°. The defocus ranges from −10 to −15 µm. For the normal dose tilt series, the total dose was 100 e^-^/Å^2^. Platelet tomograms used for model training were reconstructed using the automated pipeline in EMAN2^39^. The percentage of tilt images kept for reconstruction was 90%. The tomograms were initially output as bin2 tomograms, then another bin2 was applied by EMAN2.

### Generating tomograms with different total dosages

The high-dose tile series with a total dose of 400 e^-^/Å^2^ was achieved through increasing the exposure time of each tilt image. Each 400 e^-^/Å^2^ total dose tomogram’s micrograph was collected in 152 frames (0.052 e^-^/Å^2^/frame). All frames were corrected using MotionCor2^38^ with default settings to avoid beam-induced motion, along with the gain correction.

For the high-dose electron tomograms used for even/odd raw frames split, reconstruction was first done through EMAN2 with 100% tilt images kept on the original 400 e^-^/Å^2^ tomograms, and the alignment parameters were saved for later reduced-dose reconstruction. Micrograph raw frames used for different total doses can be found in Supplementary Table 2. The same alignment parameters were then imported with no alignment and 0 rounds of iteration using all tilt images for reduced-dose tomogram reconstruction to ensure consistency.

### Collecting data from the CryoET Data Portal

To build our dataset for training, we used API from the CryoET Data Portal (Chan Zuckerberg Imaging Institute, Chan Zuckerberg Initiative)^15^. Specifically, we searched for tomograms of sizes larger than 800*800 pixels and of magnifications smaller than 50 Å/pixel but larger than 5 Å/pixel. From these, we randomly selected 15 tomograms from each eukaryotic species and 3 tomograms from prokaryotic species that fit the selection criteria. Tomograms containing no visible biological features were excluded from our dataset after the first round of human screening. The remaining tomograms were added to our curated training dataset, as shown in Supplementary Table 1.

To generate Fig. 4c, we selectively chose both prokaryotic and eukaryotic tomograms from the cryoET tomograms generated by our lab. We limited our selection to tomograms that had an Å/pixel value of 5 or higher to match the magnification range of the tomograms used in our training set. For each magnification level, we selected 3 tomograms that, to the human eye, had obvious differences in quality and ran TomoScore to confirm these differences.

### Human quality scoring criteria

Human scoring was performed using the EMAN2^39^ 2D slice display function and its included measurement tools. Each slice of a tomogram was evaluated by eye to find the smallest recognizable feature, and then the width of that feature in pixels was measured and recorded. Pixel measurements within a range of ∼1 pixel were treated as the same measurement to reduce human error, forming five discrete categories. Features from lower-quality tomograms were more blurry and less consistent, so there was a larger variation in pixel measurements for large features. To account for the inconsistency, the ranges of categories 1 and 2 were extended to >1 pixel. A sixth category labeled “0” was included as a measurement for slices in which no features were visible.

### Sample thickness measurement

When measuring the thickness of cells using a tomogram reconstruction, we took a 2D projection along the XZ or YZ planes of the tomogram and measured the bounds of the cell by hand along the vertical Z axis (Supplementary Fig. 4a). We chose to use the YZ slices for taking measurements since the missing wedge effect leaves large artifacts along the XZ slices. Using EMAN2, we averaged 513 adjacent YZ slices to generate a 2D projection along the X-axis, smoothing out the boundaries of the cell. Measurements for cell thickness were recorded as the distance in pixels between the lower and higher cell boundaries. To ensure our measurements matched the cell boundary, we examined the Z slices we chose as the bounds for the measurement (Supplementary Fig. 4bc). Measurement bounds were reasonable if the cell was visible in the slice with no visible features, and unreasonable if the cell was not visible at all. The most recent method for sample thickness estimation by CTFFIND5^4^ makes estimations specifically for protein-level samples, while our estimation focuses more on cellular samples. Thus, this method was not included in our comparisons.

### Data preprocessing and augmentation

All tomogram slices used for this research were cropped to 960*960 pixel size and then normalized using EMAN2 before any further analysis or training. Due to the scarcity of slices with the smallest features of 4-5 pixels and 6+ pixels, rotation augmentation was applied to these slices such that slices of 6+ pixels were rotated 90 degrees 3 times (90°, 180°, 270°) and slices of 4-5 pixels were rotated 90 degrees 2 times (90°, 180°).

| Feature Diameter | No Feature | 6+ pixels | 4-5 pixels | 3 pixels | 2 pixels | 1 pixel |
| --- | --- | --- | --- | --- | --- | --- |
| #Slices after augmentation | 33,580 | 2,920 | 3,410 | 4,147 | 5,285 | 4,963 |

Gaussian filter (*σ* = 3, and *z* as the coordinate of the slice) was applied on human-judged ranks to generate continuous quality estimates across each tomogram.

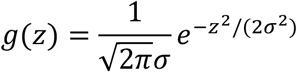

### SliceQuality model training

To achieve better generalization to ameliorate overfitting, we adopted the pre-trained weights of ResNet101 from PyTorch^40^. An adagrad variant of stochastic gradient descent was used, accompanied by mean square error loss implemented in the PyTorch with a learning rate of 5*10^−7^ for 200 epochs. Each slice of tomogram put into training was of size 960*960 pixels, and the minibatch size was 8. The model was trained on a single NVIDIA GeForce RTX 4090 GPU of 24GB of VRAM, and the model took around 72 hours to train.

## Code availability

The SliceQuality model can be downloaded from here: https://bcm.box.com/s/s418a3dhhdcpycwpmub0dzy9hsz4y58k

## Acknowledgments

We thank Hongjiang Wu and Valerie Dalton for providing suggestions on the manuscript. This work is supported by Robert Welch Foundation (X-F-0006-20240723), NIH-NHLBI (R01HL162842) to Z.W., NIH-NIAID (R01AI179879) to Z.W., NIH (R35GM151999) to S.L. We thank Tong Huo, Yuewei Wang, and Disuke Nakada for providing mouse and human platelet data. We acknowledge the use of the cryoEM supported by the Advanced Technology CPRIT Cryo-EM Core (1RP190602) at BCM and Cryo-EM core at UTHealth Center in Houston.

## Contributions

S.L. and Z.W. conceived and supervised the project. X.T. and A.A. selected and downloaded tomograms from the CryoET Data Portal^15^. Z.Y., X.Z, and X.T. performed data collection and processing. E.B. and X.T. specified scoring criteria. E.B. and A.A. scored all the tomograms. X.T. performed model training and evaluation. X.T., S.L., and Z.W. wrote the manuscript with other authors’ input.

## Notes

### Competing Interest Statement

The authors have declared no competing interest.

### Summary of Updates

1. Model Robustness: We revised the manuscript to better describe the robustness of the TomoScore model across a wide range of tomograms and imaging conditions. The previous mention of optimizing electron dose has been removed to clarify the scope of our analysis. 2. Sample Thickness Validation: We now include a quantitative assessment and validation of sample thickness, with a new Supplementary Figure (Supplementary Fig. 6) that illustrates this metric across different datasets. 3. Expanded Test Set: To strengthen our validation, we increased the size of the test dataset. Figure 2 now includes a gallery of representative tomograms spanning a range of TomoScore values to demonstrate the model performance in diverse conditions. 4. Performance Benchmarking: We added benchmarking data on model inference time to highlight the practical utility of TomoScore in high-throughput workflows. 5. Clarified Abstract and Introduction: We rephrased key sections of the abstract and introduction to address and prevent misunderstandings raised by reviewers, particularly regarding the model scope, capabilities, and intended applications. 6. Revised Supplementary Figures: Supplementary Figures 1 and 2 have been updated to more clearly illustrate the criteria and consistency of human scoring used during training and validation. 7. Author order modified.

