## supplements for "TomoScore: A Neural Network Approach for Quality Assessment of Cellular cryoET"

### Supplementary information

| | Species | #Tomograms | #Slices |
|---|---|---|---|
| **Eukaryote** | Chlamydomonas reinhardtii | 10 | 4616 |
| | Homo sapiens | 2 | 981 |
| | Homo sapiens (T-cell) | 13 | 3632 |
| | Homo sapiens (HUVEC) | 5 | 2150 |
| | Schizosaccharomyces pombe | 8 | 4972 |
| | Mus musculus | 5 | 1419 |
| | Rattus rattus (pancreatic beta cells) | 10 | 6078 |
| **Prokaryote** | Bacillus subtilis | 3 | 1126 |
| | Escherichia coli | 2 | 1791 |
| | Helicobacter pylori | 2 | 1293 |
| | Legionella pneumophila | 3 | 1489 |
| | Helicobacter hepaticus | 3 | 2388 |
| | Magnetospirillum magneticum | 1 | 700 |
| | Nitrosopumilus maritimus | 3 | 1488 |
| | Pseudomonas aeruginosa | 3 | 1789 |
| | Spiroplasma melliferum | 1 | 157 |
| | Streptococcus pneumoniae | 3 | 2092 |
| | Sulfolobus acidocaldarius | 3 | 2091 |
| | Sulfolobus solfataricus | 2 | 1400 |
| | Treponema primitia | 1 | 400 |
| | Vibrio cholerae | 5 | 4587 |
| **Anucleate Eukaryote (Platelets)** | Homo sapiens | 9 | 3395 |
| | Mus musculus | 17 | 8492 |
| **Total** | **19** | **114** | **58527** |

**Table 1 Summary of tomogram dataset**

| TOTAL DOSAGE | FRAMES/ANGLE | SELECTED FRAMES (EVEN/ODD) |
|---|---|---|
| 400 $e^-/Å^2$ | 152 | #1, #2, #3, #4, …, #152 |
| 200 $e^-/Å^2$ | 76 | #2, #4, #6, #8, …, #152 / #1, #3, #5, #7, …, #151 |
| 100 $e^-/Å^2$ | 38 | #2, #6, #8, #12, …, #152 / #1, #5, #7, #11, …, #151 |
| 50 $e^-/Å^2$ | 19 | #2, #10, #18, #26, …, #152 / #1, #9, #17, #25, …, #151 |
| 25 $e^-/Å^2$ | 9 | #10, #26, #42, #58, … #138 / #9, ##25, #41, #57, …, #137 |
| 13 $e^-/Å^2$ | 5 | #10, #40, #70, #100, #130 / #9, #39, #69, #99, #129 |

**Table 2 Selected raw frames for different total dosage tomograms' tilt series.**

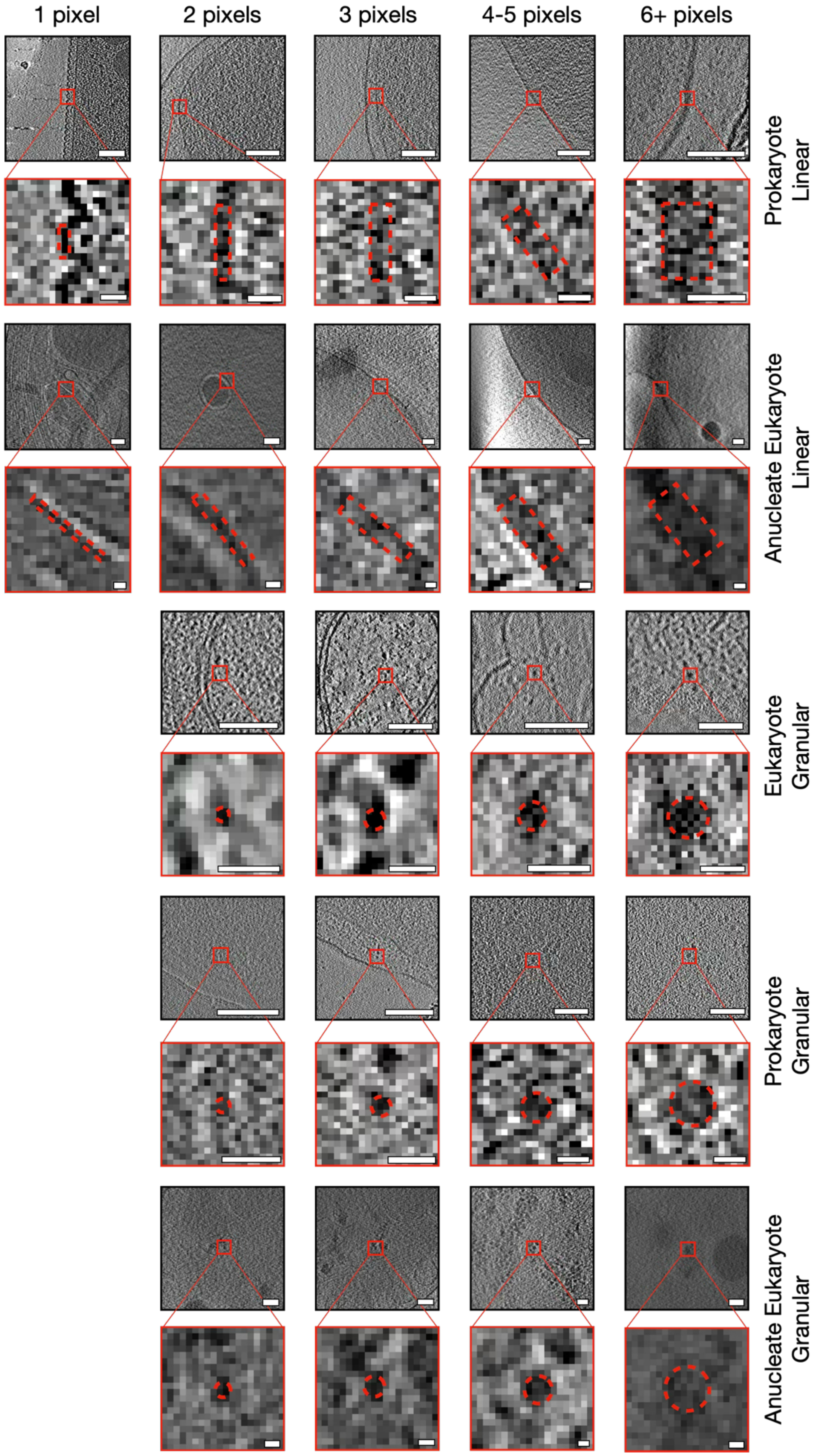
1 pixel
2 pixels
3 pixels
4-5 pixels
6+ pixels
Prokaryote Linear
Anucleate Eukaryote Linear
Eukaryote Granular
Prokaryote Granular
Anucleate Eukaryote Granular

**Supplementary Figure 1 Tomogram slices with quality-determining features**

Additional examples of feature measurements in Eukaryotes (top rows), Prokaryotes (middle rows), and Anucleate Eukaryotes (bottom rows). Features are separated into membrane structures (left group), and particle structures (right group). Membrane features are measured by their width, while particle features are measured by their diameter. Black-bounded images are cropped to 200x200 pixels and have a scale bar of 100 nm. Red-bounded images are cropped to 20x20 pixels and have a scale bar of 10 nm.

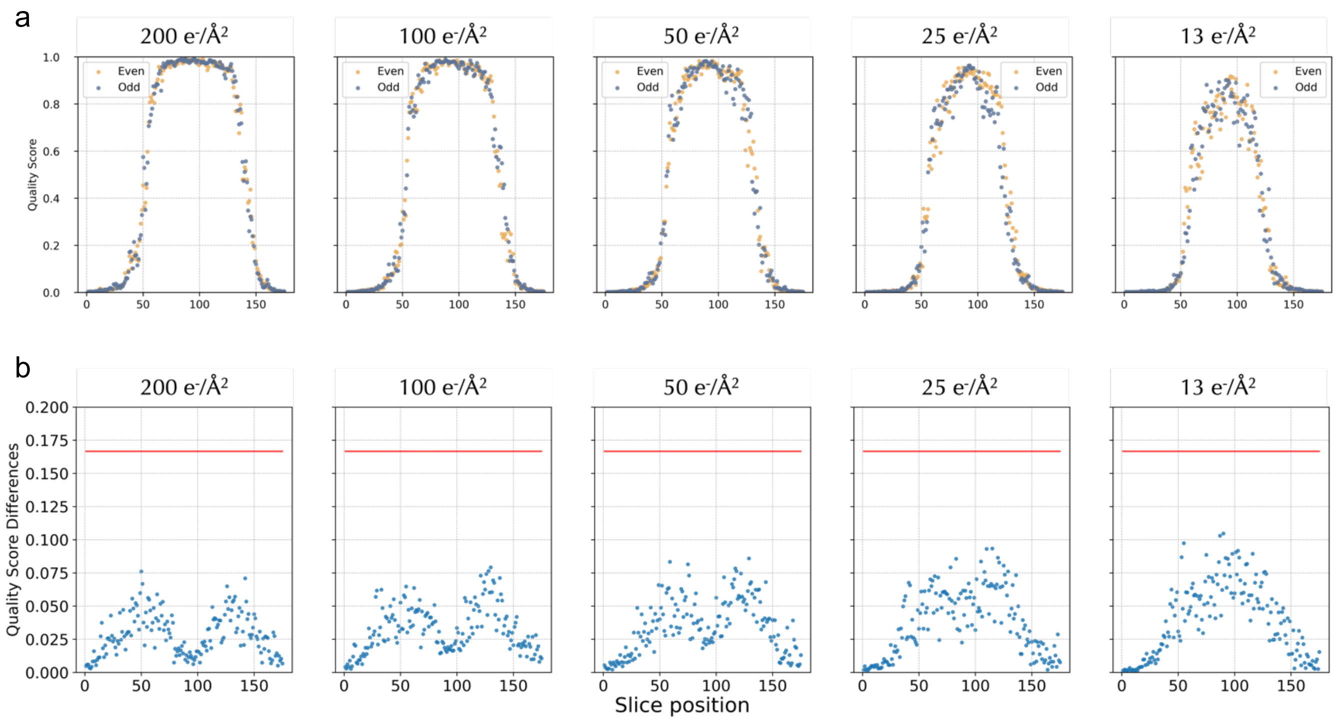


**Supplementary Figure 2 Even/Odd frames reconstructed tomograms are "identical"**

a) Shows the SliceQuality Network's predictions for the same set of tomograms, where the quality scores predicted for even/odd splits of different decreasing total dosages. b) Average quality score differences of 10 tomograms split into different dosages vary within a small range, indicating the high consistency of our trained model.

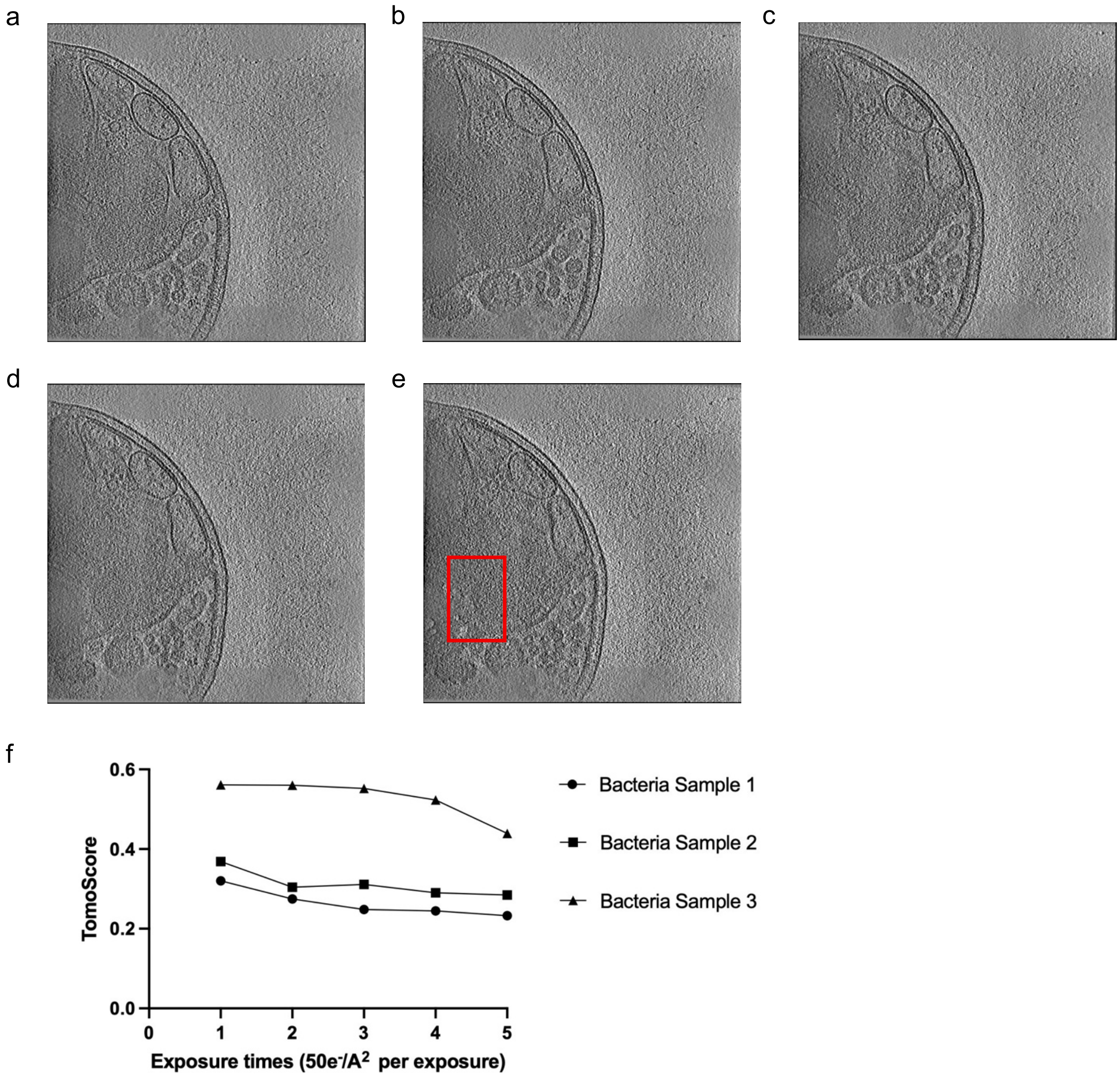


**Supplementary Figure 3 TomoScore decreases with increasing electron dose in E. coli.**

a) - e) Representative tomogram slices for E. coli with different cumulative exposure doses from 50e-/A$^2$ to 250e-/A$^2$. The red box indicates the radiation bubbles. f) TomoScore values for two independent E. coli samples decrease as the cumulative electron dose increases from 50 to 250 $e^-/Å^2$.

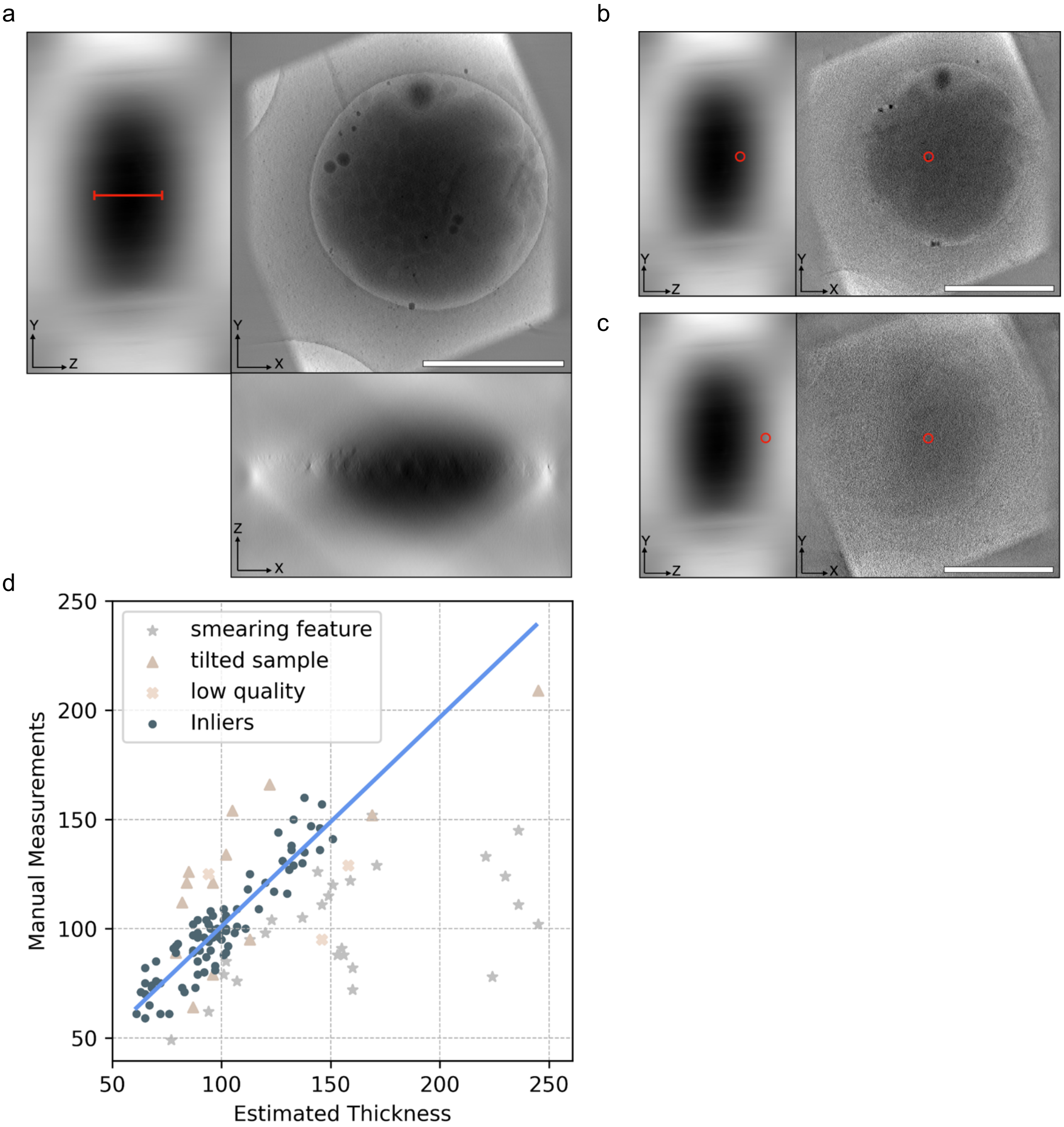


**Supplementary Figure 4 Sample thickness estimation and validation**

a) YZ, XY, and XZ projections of a platelet tomogram. The red bar shows which part of the image was used to measure cell thickness. b) The red circle indicates a point on the measured edge of the cell on the YZ projection. c) The red circle indicates a suggested alternative point of measurement taken at a lower contrast boundary. All scale bars are 1000 nm. d) RANSAC regression pinpointed inliers and outliers of the thickness estimations. 137 tomograms tested for thickness were all platelet cells. Reconstruction thickness varied from 175 to 600 slices in the z-direction.
